# Role of α-tubulin helix 11’ in heterodimer conformation and microtubule dynamics

**DOI:** 10.64898/2026.02.16.705813

**Authors:** Katelyn J. Hoff, Kaitlin Alemany, Jeffrey K. Moore

## Abstract

Tubulin heterodimers transition from a curved conformation in solution to straight conformation when assembled into the microtubule lattice. Many proteins and small molecules alter microtubule dynamics by binding and stabilizing the curved or straight conformations; however, we have a poor understanding of the regions of the tubulin heterodimer that contribute to this transition and whether these conformations represent sources of phenotypic variation in human disease and across species. We previously identified a role for α-tubulin helix 11’ (H11’) in the curved-to-straight transition. Using computational simulations and tubulin mutants in budding yeast, we show that H11’ variants associated with human disease destabilize the curved conformation of the heterodimer and disrupt α-tubulin function in cells. Comparing α-tubulin sequences across eukaryotes demonstrates strong conservation of H11’ with the exception of several α-tubulin isotypes expressed during mitosis in the amoebae *Naegleria fowleri* and *Naegleria gruberi*. Introducing H11’ sequence variants from *Naegleria* in budding yeast α-tubulin increases heterodimer exchange at microtubule plus ends and destabilizes mitotic spindles. We provide evidence that interactions between α-tubulin H11’ and β-tubulin H8 stabilize the curved conformation of tubulin heterodimers, and that the equilibrium between curved and straight conformations is an ancient feature of tubulin evolution.

## Introduction

Microtubules are composed of α– and β-tubulin heterodimers that bind to growing ends during states of polymerization and rapidly unbind during depolymerization, a unique process known as “dynamic instability”^1^. Although dynamic instability is an intrinsic property of tubulins, organisms have evolved diverse regulatory mechanisms to finely-tune microtubule dynamics in different contexts^2^. These regulatory mechanisms control processes including chromosome segregation during cell division, cargo transport across long-range cell processes such as axons, and a variety of other key biological processes across eukaryotes. Therefore, understanding the fundamental control points that these regulatory mechanisms act upon is essential for our understanding of the evolution of microtubule-based processes and how these may be disrupted in human disease.

Microtubule dynamics are regulated intrinsically, through the interactions and conformational dynamics of tubulin proteins, and extrinsically, through microtubule-associated proteins (MAPs) and motors that bind to and influence tubulin conformation. Intrinsic regulation is controlled by the GTP-binding and hydrolysis cycle of the tubulin heterodimer, which promote the axial compaction and twisting of the αβ-heterodimer^3–9^. In addition, heterodimers adopt curved or straight conformational states that are not influenced by nucleotide state, but are important for microtubule polymerization and depolymerization^10–15^. Heterodimers can alternate between the curved and straight conformations in solution or at microtubule ends, but are trapped in the straight conformation when bound by neighboring tubulins in the microtubule lattice. Conformational dynamics of tubulin heterodimers can vary across species, and are thought to create inter-species differences in microtubule dynamics^16–19^. In addition, missense variants in α– and β-tubulin isotypes identified in human neurodevelopmental disorders known as ‘tubulinopathies’ have been shown to impact tubulin activity by altering nucleotide-dependent^20^ and nucleotide-independent conformational changes^21–24^. Importantly, neither extrinsic nor intrinsic processes work in isolation, and a major goal of the field is to understand how intrinsic and extrinsic regulation synergize to control microtubule dynamics in cells.

Our previous work showed that tubulinopathy missense variants which create alanine and isoleucine substitutions at residue 409 in α-tubulin (V409A and V409I, respectively) dominantly alter tubulin’s intrinsic assembly activity and subvert normal regulation conferred by XMAP215/Stu2^23^. XMAP215/Stu2 proteins regulate microtubule dynamics by combining plus-end localization activity with binding to αβ-heterodimers in the curved conformation^25–28^. Residue 409 of α-tubulin is located in helix 11’ (H11’) near the interface between α– and β-tubulin in the heterodimer and serves as a hinge point between the curved and straight heterodimer conformations. We proposed a model in which these residue 409 variants destabilize the curved conformation of the tubulin heterodimer and accordingly promote a predominantly straight conformation that is activated for microtubule assembly^23^.

In this study we investigate the role of α-tubulin helix 11’ (H11’) in tubulin conformation and microtubule dynamics. We investigate the molecular function of α-tubulin H11’ using computational tools to predict the conservation of α-tubulin secondary structure, as well as the effects of H11’ mutations on overall tubulin conformational stability. We find that α-tubulin H11’ is highly conserved across species, with a notable exception in species of the amoeba *Naegleria*. These *Naegleria* species express α-tubulin isotypes during mitosis that harbor amino acid sequence variants in α-tubulin H11’. By modeling these in budding yeast α-tubulin, we find that the H11’ variants increase rates of tubulin assembly and disassembly, and disrupt the stability of the mitotic spindle. Together, these findings demonstrate an important role for α-tubulin H11’ in the conformational dynamics and assembly activity of tubulin, and suggest that H11’ may act as a control point for dynamic instability across evolution.

## Results

### α-tubulin helix 11’ variants are predicted to disrupt the stability of the curved tubulin conformation

We used the DUET server^29^ to predict the thermodynamic effects of residue substitutions at V409 on the curved heterodimer, as represented in a structure of *Sus scrofa* TUBA1A and TUBB α/β-tubulin heterodimer bound to the TOG1 and TOG2 domains of the fission yeast XMAP215, Alp14 (PDB:6MZG;^28^; Figure 1A). In these simulations, an increased ΔΔG value indicates that the variant destabilizes the protein structure^29^. Among the 19 alternative amino acid substitutions at α-V409, the V409I tubulinopathy variant is predicted to have a mildly destabilizing effect (0.896 kcal/mol), while the V409A tubulinopathy variant is predicted to be twice as destabilizing as V409I (1.886 kcal/mol) (Figure 1B). In sampling all possible substitutions at this position, we found that V409D, although not a tubulinopathy variant previously observed in patients, is predicted to have the strongest destabilizing effect (2.656 kcal/mol) (Figure 1B). These data suggest that the stability of the curved heterodimer conformation is sensitive to amino acid substitutions at position 409, and that the magnitude of impact depends on the side chain of the substituted amino acid.

**Figure 1.**
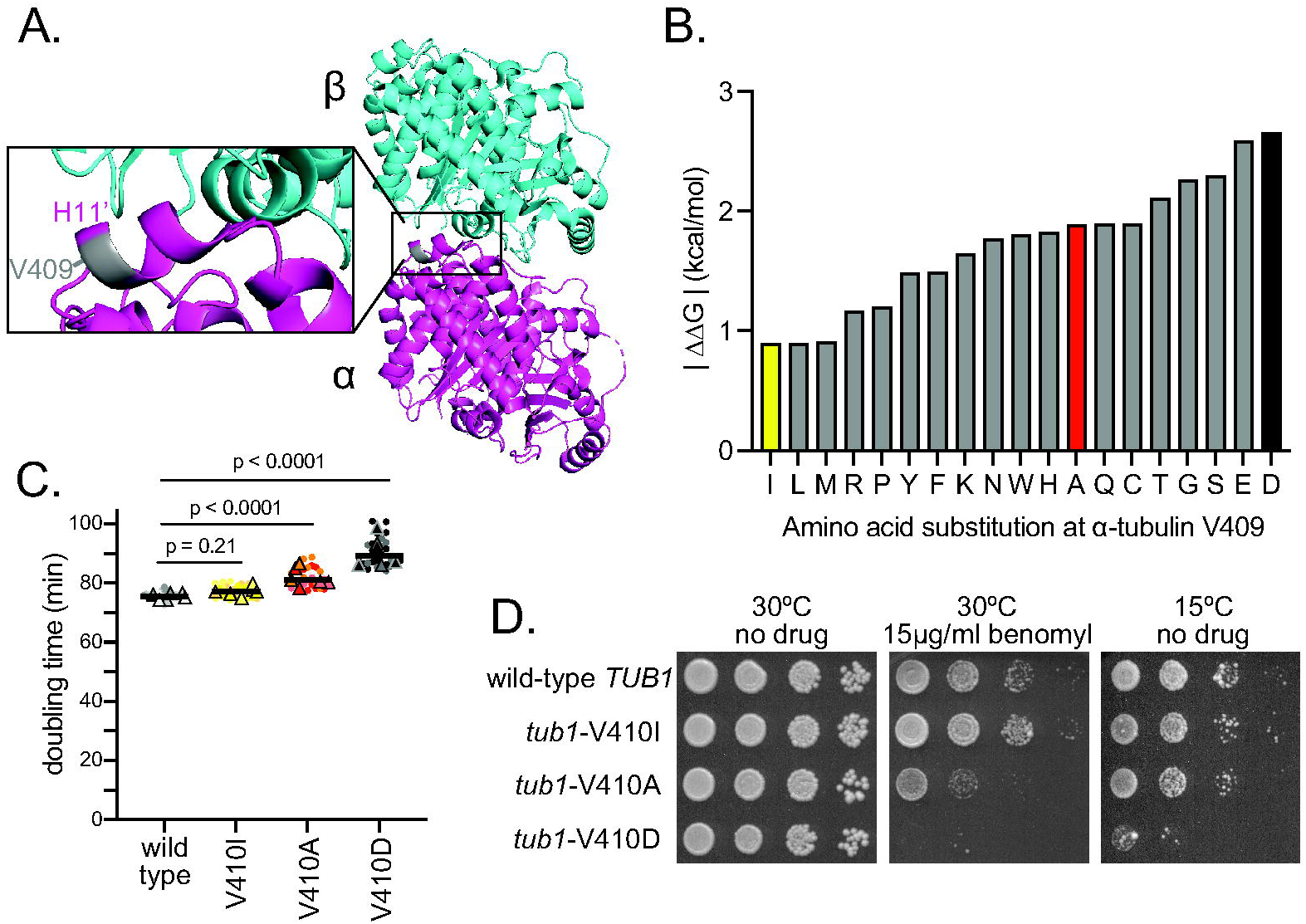
Tubulinopathy variants in α-tubulin helix 11’ disrupt the curved tubulin conformation. (**A**) *Sus scrofa* TUBA1A/TUBB heterodimer structure (PDB:6MZG)^28^ with α-tubulin in magenta and β-tubulin in cyan. Inset highlights residue Valine 409 in grey, which resides in α-tubulin H11’. **(B)** DUET predicted ΔΔG values for the V409 residue when replaced by any of the other 19 amino acids. Tubulinopathy variants V409I and V409A highlighted in yellow and red, respectively. V409D, highlighted in black, has not been identified in patients or associated with disease, but is the variant predicted to be the most destabilizing to the curved tubulin conformation. **(C)** Doubling times of cells expressing the indicated mutation at the genomic *TUB1* locus. Dots represent technical replicates, colors indicate biological replicates and triangles indicate mean values of six independent experiments. Six technical replicates were used in each experiment, and two biological replicates were used per genotype across the independent experiments. Lines represent median value across replicates. Statistical analysis was done using an ordinary one-way ANOVA analyzed post hoc by a Tukey test. **(D)** Tenfold dilution series of the indicated strains were spotted to rich medium or rich medium supplemented with the indicated concentration of benomyl. Cells were grown at 30°C for two days or 15°C for twelve days.

We next sought to confirm the computational predictions in a biological model of tubulin function. Accordingly, we made three mutations in the budding yeast α-tubulin (*TUB1*) at position V410, which corresponds to residue V409 in mammalian TUBA1A. We constructed *tub1*-V410I and –V410A, which represent the two tubulinopathy variants, as well as *tub1*-V410D to represent the most severe destabilizing effect on the curved conformation (Figure 1B). Each of these mutant strains is viable as a haploid expressing the V410 mutant at the native *TUB1* locus and a wild-type allele of the minor α-tubulin *TUB3* at its native locus. To assess tubulin function, we first measured the doubling times of each mutant strain compared to wild-type controls. Mutants that disrupt tubulin function in yeast are known to cause errors in mitosis, and the spindle assembly checkpoint prolongs mitosis until these errors are corrected^30, 31^. Compared to wild type, we find that *tub1*-V410I has no significant change in doubling time, while *tub1*-V410A exhibits an approximately 8.5% increase (p < 0.0001, compared to WT; Figure 1C; Table 1). *tub1*-V410D cells exhibit a more severe doubling time increase of 20% (p < 0.0001, compared to WT; Figure 1C). These results indicate that the proliferation rates of the *tub1*-V410 mutant cells scale with the severity of destabilizing effects predicted by computational modeling.

**Table 1.** Doubling times for H11’ mutants in budding yeast. Values are mean minutes ± SD. Boldface indicates p < 0.01, based on t-test vs wild-type control in the same media condition.

| α-tubulin/TUB1 allele | Doubling time in rich media | Doubling time in 1% DMSO | Doubling time in 1.25 μM nocodazole, 1% DMSO |
| --- | --- | --- | --- |
| TUB1 (WT) | 78 ± 5 | 94 ± 4 | 99 ± 4 |
| V410A | **85 ± 7** | nd | nd |
| V410I | 77 ± 2 | nd | nd |
| V410D | **91 ± 5** | nd | nd |
| F405Y | 83 ± 4 | 97 ± 4 | **450 ± 10** |
| W408H | **86 ± 4** | 94 ± 3 | 99 ± 4 |
| Y409F | 78 ± 3 | 91 ± 5 | 95 ± 6 |
| F405Y, W408H, Y409F (YHF) | **85 ± 6** | 97 ± 6 | **461 ± 32** |

Next, we confirmed the predicted destabilizing effects by assaying sensitivity to other stimuli that disrupt the conformational dynamics of αβ-heterodimers, predicting that these might exhibit additive effects for V410 mutants. We first tested sensitivity to benomyl, a drug that binds tubulin and prevents microtubule assembly. Although the structure of the benomyl-bound state of tubulin has not been solved, it is not thought to represent the curved conformation since benomyl does not interfere with the binding or activity of colchicine^32^. We also tested sensitivity to low temperature, which slows microtubule assembly and disassembly and inhibits the recycling of soluble αβ-heterodimers into an assembly competent state^33^. The *tub1*-V410I mutant cells exhibit a level of benomyl sensitivity and low temperature sensitivity that is similar to wild type (Figure 1D). In contrast, *tub1*-V410A mutant cells are hypersensitive to benomyl but are not obviously more sensitive to low temperature than wild-type controls, while the *tub1*-V410D mutant cells are hypersensitive to both benomyl and low temperature (Figure 1D). Together, the data from our proliferation and microtubule stability assays support the predictions from computational modeling, with the cellular phenotype scaling with the predicted effect on protein stability. These results suggest that α-tubulin H11’ normally stabilizes the curved heterodimer conformation.

### **α**-tubulin helix 11’ is highly conserved across species and tubulin isotypes

If H11’ stabilizes the curved conformation of tubulin, we reasoned that this domain should be strongly conserved across species and tubulin isotypes. We compared the conservation of H11’ to all other secondary structure features of α-tubulin by aligning 1,347 α-tubulin sequences from 438 species across the eukaryotic tree of life^34, 35^. We then used this multiple sequence alignment to determine conservation scores using the ConSurf server^36,37^. The continuous conservation scores calculated for each residue were then binned into a discrete scale ranging from highly variable (grade 1; represented in green) to highly conserved (grade 9; represented in purple), and the conservation grade classification for each residue was mapped onto a structure of *Sus scrofa* TUBA1A (Figure 2A; PDB: 6MZG;^28^). As an alternative visualization, we calculated the percentage of the 1,347 α-tubulins that retain the consensus amino acid at each position and binned the data by α-tubulin secondary structure features. We excluded unstructured loops from this analysis since these tend to exhibit more variability than α-helices and β-sheets. We find that amongst all secondary structures, α-tubulin H11’ is the most highly-conserved secondary structure feature with the lowest degree of variation (96.6%±1.8; Table 2; Figure 2B). These data indicate that while many regions of α-tubulin are highly conserved across eukaryotes, H11’ exhibits the strongest conservation.

**Figure 2.**
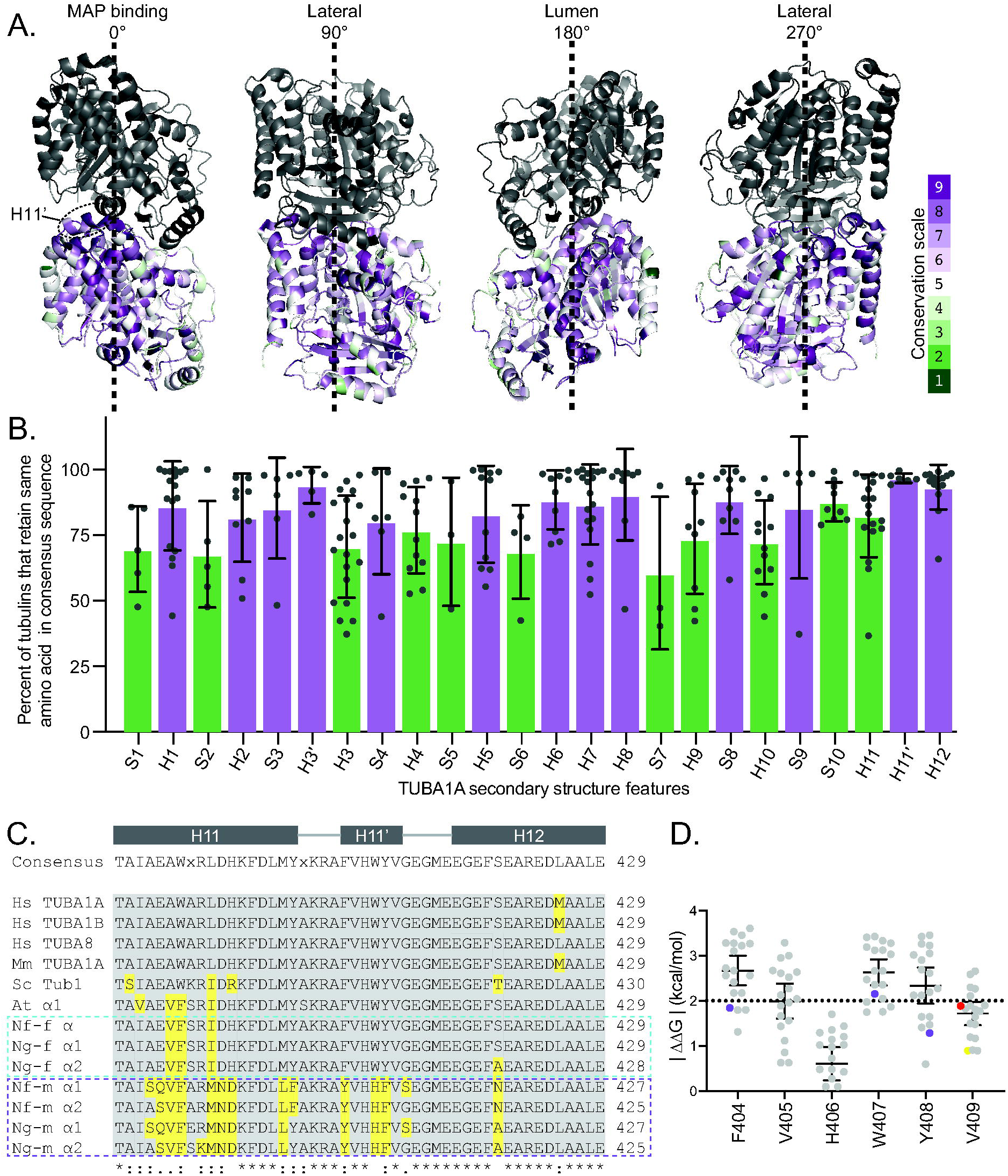
α-tubulin helix 11’ is highly conserved across species and tubulin isotypes. **(A)** Mapping the conservation score of each residue on α-tubulin calculated by the ConSurf server^36,37^. The scores were mapped onto a *Sus scrofa* TUBA1A/TUBB heterodimer structure (PDB:6MZG)^28^. The conservation scores range from variable (1, indicated as green) to conserved (9, indicated as purple). Four views of the heterodimer are shown: 0° represents the MAP binding surface, 90° and 270° represent lateral interfaces, and 180° represents the luminal-facing side of the microtubule. **(B)** The percentage of the 1,347 α-tubulin sequences^34^ analyzed that retain the amino acid observed in the consensus sequence that was determined from the multiple sequence alignment of this same data set. Each dot represents an individual residue within the indicated secondary structure of TUBA1A. Bars represent mean value of each data point and error bars represent standard deviation. The mean value of each secondary structure was statistically compared to H11’ using an unpaired t-test. **(C)** Sequence alignment of α-tubulin helix 11, helix 11’, and helix 12 from a representative list of eukaryotes generated by Clustal Omega^55,56,57^. Consensus sequence generated from aligning the previously described 1,347 α-tubulin sequences. Grey and yellow backgrounds represent a conserved or divergent residue, respectively, when compared to the consensus sequence. **(D)** Plot represents the change in the change of Gibbs free energy (ΔΔG) values predicted by the DUET server^29^ when the indicated residue is mutated to any of the other 19 amino acids. These predicted values are based on importing the *Sus scrofa* TUBA1A/TUBB heterodimer bound to Alp14 TOG1 and TOG2 domains (PDB:6MZG)^28^. Purple dots represent variants identified at these residues in *Naegleria* mitotic α-tubulins. Red and yellow dots represent V409A and V409I patient mutations, respectively. Dotted line at 2kcal/mol represents the minimum ΔΔG value that is predicted to have a highly destabilizing effect on the imported protein conformation structure. Lines and error bars represent mean value ± 95% confidence interval.

**Table 2.** Conservation of amino acid sequence across α-tubulin secondary structure features. Values are average conservation of the consensus residues within the secondary structure feature. Boldface indicates p-value ≤ 0.05 in t-test comparing to H11’.

| TUBA1A secondary structure | 2° Structure Feature | Mean Percentage ± Standard Deviation |
| --- | --- | --- |
|  | **S1** | **69.6 ± 16.3** |
| | H1 | 86.1 ± 16.9 |
|  | **S2** | **67.7 ± 20.3** |
| | H2 | 81.6 ± 16.8 |
| | S3 | 85.2 ± 19.2 |
| | H3' | 94.0 ± 6.9 |
|  | **H3** | **70.6 ± 19.4** |
| | S4 | 80.2 ± 20.2 |
|  | **H4** | **76.8 ± 16.4** |
|  | **S5** | **72.5 ± 24.4** |
| | H5 | 82.9 ± 18.3 |
|  | **S6** | **68.6 ± 17.8** |
| | H6 | 88.4 ± 11.3 |
| | H7 | 86.6 ± 15.2 |
| | H8 | 90.4 ± 17.4 |
|  | **S7** | **60.5 ± 29.1** |
|  | **H9** | **73.6 ± 21.0** |
| | S8 | 88.3 ± 12.9 |
|  | **H10** | **72.3 ± 15.9** |
| | S9 | 85.5 ± 27.0 |
|  | **S10** | **87.6 ± 7.4** |
|  | **H11** | **82.3 ± 15.8** |
| | H11' | 96.6 ± 1.8 |
| | H12 | 93.3 ± 8.5 |

### Helix 11’ variants in *Naegleria* alter α-tubulin function

While our analysis shows strong conservation of H11’ across species, we did identify variants in the α-tubulins of amoebae *Naegleria fowleri* and *Naegleria gruberi*. These amoebae express two distinct sets of α– and β-tubulin isotypes, one to support the assembly and maintenance of flagella (Figure 2C, cyan outline) and another to support mitotic cell division (Figure 2C, purple outline)^38,35^. The α-tubulins expressed in the flagellate state of both species have a H11’ that is identical to the sequence in other eukaryotes. However, the α-tubulins expressed in the mitotic state include three divergent residues in H11’ – at positions 404, 407, and 408 (Figure 2C). These mitotic α-tubulins also exhibit sequence divergence in neighboring regions such as helix 11 and helix 12; however, this divergence was also observed in the flagellar α-tubulins and was not consistently divergent between the two *Naegleria* species (Figure 2C). Therefore, we focused on the three divergent residues observed in H11’ in the *Naegleria* mitotic α-tubulins.

We next asked whether the variant residues identified in the *Naegleria* mitotic α-tubulin H11’ impact tubulin’s curved conformation stability, using the same computational modeling approach as applied to the tubulinopathy variants at V409 (Figure 1). Changing each of the consensus F404, W407, and Y408 residues individually to any of the other 19 amino acids creates mean ΔΔG values greater than two, indicating that variants at these positions are highly destabilizing (Figure 2D;^29^). The amino acid identities from *Naegleria* mitotic α-tubulins, F404Y, W407H, and Y408F are predicted to create milder destabilizing effects than other potential substitutions at those positions (1.841, 2.153, 1.284 kcal/mol, respectively; highlighted in purple in Figure 2D). These values are comparable to the ΔΔG values and relative distributions of the tubulinopathy variants, V409I and V409A (highlighted in yellow and red, respectively, in Figure 2D), suggesting that the *Naegleria* H11’ variants may destabilize the curved heterodimer conformation to a degree similar to the tubulinopathy variants.

To study the impact of the divergent residues in *Naegleria* mitotic α-tubulin H11’ and uncover the functional roles of this helix, we mutated the corresponding residues in budding yeast individually (referred to as *tub1*-F405Y, *tub1*-W408H, or *tub1*-Y409F), or in combination (referred to as *tub1*-YHF; Figure 3A). First, we tested whether any of these mutations prevent α-tubulin assembly into yeast microtubules by fusing each mutant α-tubulin to GFP and expressing them ectopically in yeast that also express untagged, wild-type α-tubulin. Each mutant-GFP fusion localized to microtubules, indicating that the divergent residues in *Naegleria* mitotic α-tubulin H11’ do not prevent α-tubulin assembly into yeast microtubules (Figure 3B).

**Figure 3.**
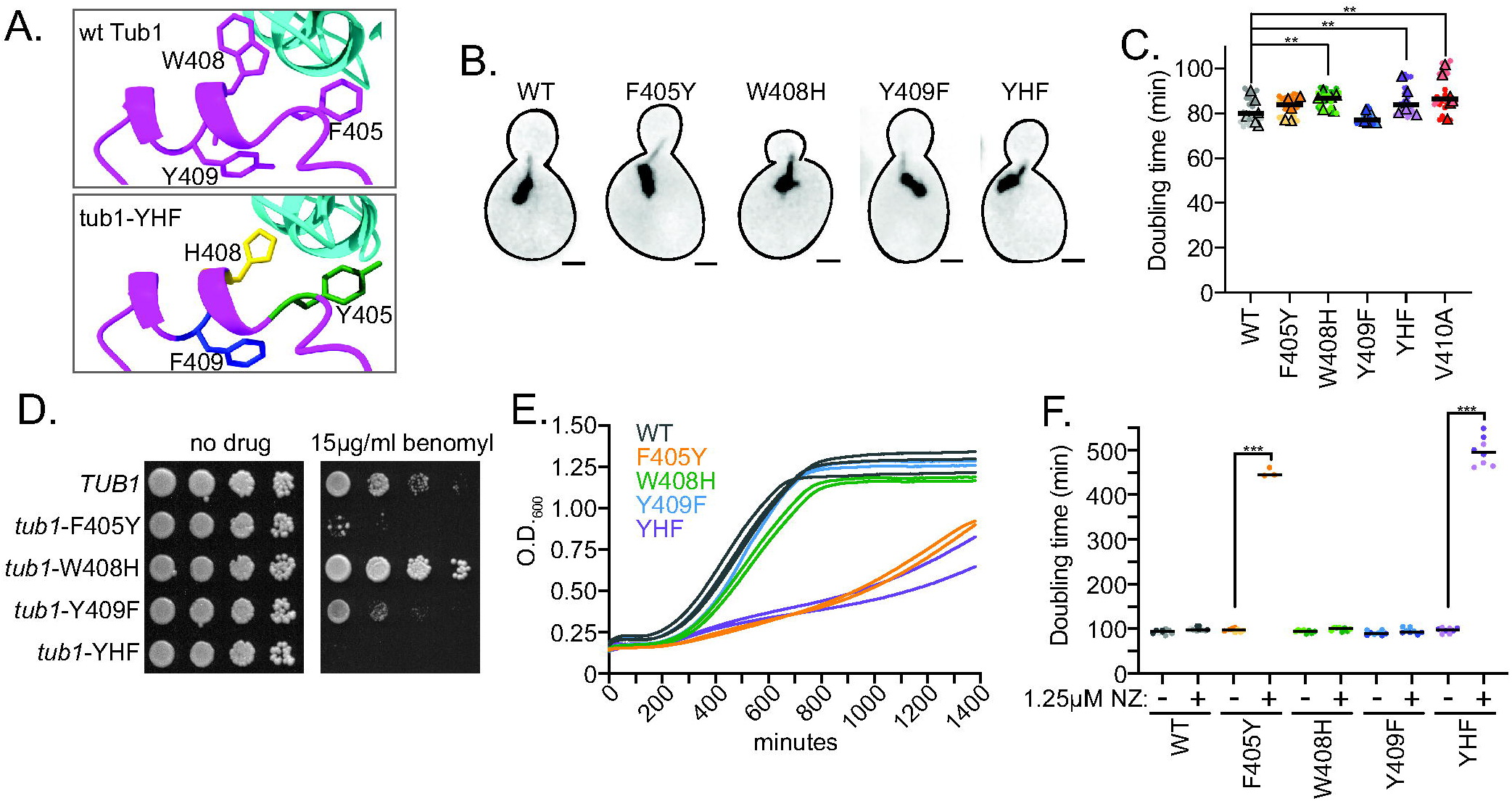
*Naegleria* α-tubulin helix 11’ variants disrupt budding yeast fitness and sensitivity to microtubule destabilizers. **(A)** Model of budding yeast α/Tub1 (magenta) and β/Tub2 (cyan) heterodimer showing sidechains of the wild-type residues F405, W408 and Y409 or the variant residues Y405, H408 and F409 in H11’. Generated using AlphaFold3^62^. **(B)** Representative images of cells expressing the indicated mutant integrated at the genomic *TUB1* locus, along with GFP-Tub1 that is either WT or the mutant with the same mutation. At least 30 cells were imaged per genotype. Scale bar = 1μm. **(C)** Doubling times of cells expressing the indicated mutation at the genomic *TUB1* locus. Dots represent technical replicates, colors indicate biological replicates and triangles indicate mean values of six independent experiments. Six technical replicates were used in each experiment, and two biological replicates were used per genotype across the independent experiments. Lines represent median values from all replicates. Asterisks indicate p < 0.001 based on Mann-Whitney test comparing to wild-type controls. **(D)** Ten-fold dilution series of the indicated strains were spotted to rich medium or rich medium supplemented with the indicated concentration of benomyl. Cells were grown at 30°C for two days. **(E)** Example growth curves of wild-type controls and indicated tub1 mutants in the presence of 1.25 µM nocodazole. Colored lines indicate biological replicates of the indicated genotype. **(F)** Doubling times of cells expressing the indicated mutation at the genomic *TUB1* locus, either in 1% DMSO or 1% DMSO with 1.25 µM nocodazole. Dots represent technical replicates and colors indicate biological replicates. Three technical replicates and two biological replicates were used per genotype in each experiment, and three independent experiments were performed. Lines represent median value across replicates. Asterisks indicate p < 0.0001 based on Mann-Whitney test comparing DMSO control and 1.25 µM nocodazole.

We then integrated each mutant into the native *TUB1* locus, replacing wild-type *TUB1*, and tested for rescue of α-tubulin function. Haploid cells expressing each mutant as the only copy of *TUB1* are viable, but exhibit different effects on proliferation and/or sensitivity to microtubule destabilizing drugs. In the proliferation assay, *tub1*-W408H single mutant and *tub1*-YHF triple mutant exhibit a 9% and 5% increase in doubling time, respectively, while the *tub1*-F405Y and *tub1*-Y409F single mutants proliferate similarly to wild-type controls (Figure 3C; Table 1). These results indicate that the W408H substitution alone and the combination of F405Y, W408H, andY409F in the YHF triple mutant disrupt α-tubulin function during mitosis.

We used two experiments to test the sensitivity of each mutant to destabilizing stimuli. In the benomyl sensitivity assay, the *tub1*-F405Y and *tub1*-Y409F single mutants are more sensitive to benomyl than wild-type controls, while the *tub1*-W408H single mutant is resistant to benomyl (Figure 3D). The *tub1*-YHF triple mutant is hypersensitive to benomyl, with a stronger phenotype than either *tub1*-F405Y or *tub1*-Y409F single mutants (Figure 3D). To determine whether these phenotypes were specific to benomyl, or generalizable to other depolymerizing stimuli, we measured sensitivity to nocodazole, a drug that inhibits polymerization by binding β-tubulin and stabilizing the curved conformation of the heterodimer^39^. For these experiments, we measured the proliferation of cells in liquid culture supplemented with 1% DMSO to maintain the solubility of nocodazole. Both the *tub1*-F405Y single mutant and *tub1*-YHF triple mutant exhibit increased sensitivity to low levels of nocodazole (1.25 µM; Figure 3E and F; Table 1). In contrast, the *tub1*-W408H and *tub1*-Y409F single mutants are similar to wild-type controls, even at higher concentrations of nocodazole (Figure 3E and F; data not shown). Together, these results indicate that the F405Y and W408H mutations individually alter α-tubulin function and combining these mutations with *tub1*-Y409F, as in *tub1*-YHF and in the *Naegleria* mitotic α-tubulins, alters α-tubulin function in a way that is distinct from any single mutant alone.

### **α**-tubulin helix 11’ alters heterodimer exchange at the microtubule plus end

We next sought to determine how the divergent H11’ of *Naegleria* mitotic α-tubulin impacts microtubule dynamics. We focused on the YHF triple mutant and W408H single mutant, since these have strong but opposing effects on benomyl sensitivity (Figure 3D). To measure microtubule dynamics, we tagged the microtubule plus-end tracking protein Bik1/CLIP-170 at its native locus with three copies of GFP (Figure 4A). We then imaged living cells at four-second intervals for ten minutes and measured changes in microtubule length over time. Figure 4B shows example traces of single microtubules from wild-type, *tub1*-W408H, and *tub1-*YHF cells. Our cumulative measurements show that *tub1*-W408H and *tub1-*YHF cells exhibit longer astral microtubules than wild-type controls (Figure 4C; median lengths per cell, 0.52, 0.97, 1.14 µm for WT, W408H, and YHF, respectively).

**Figure 4.**
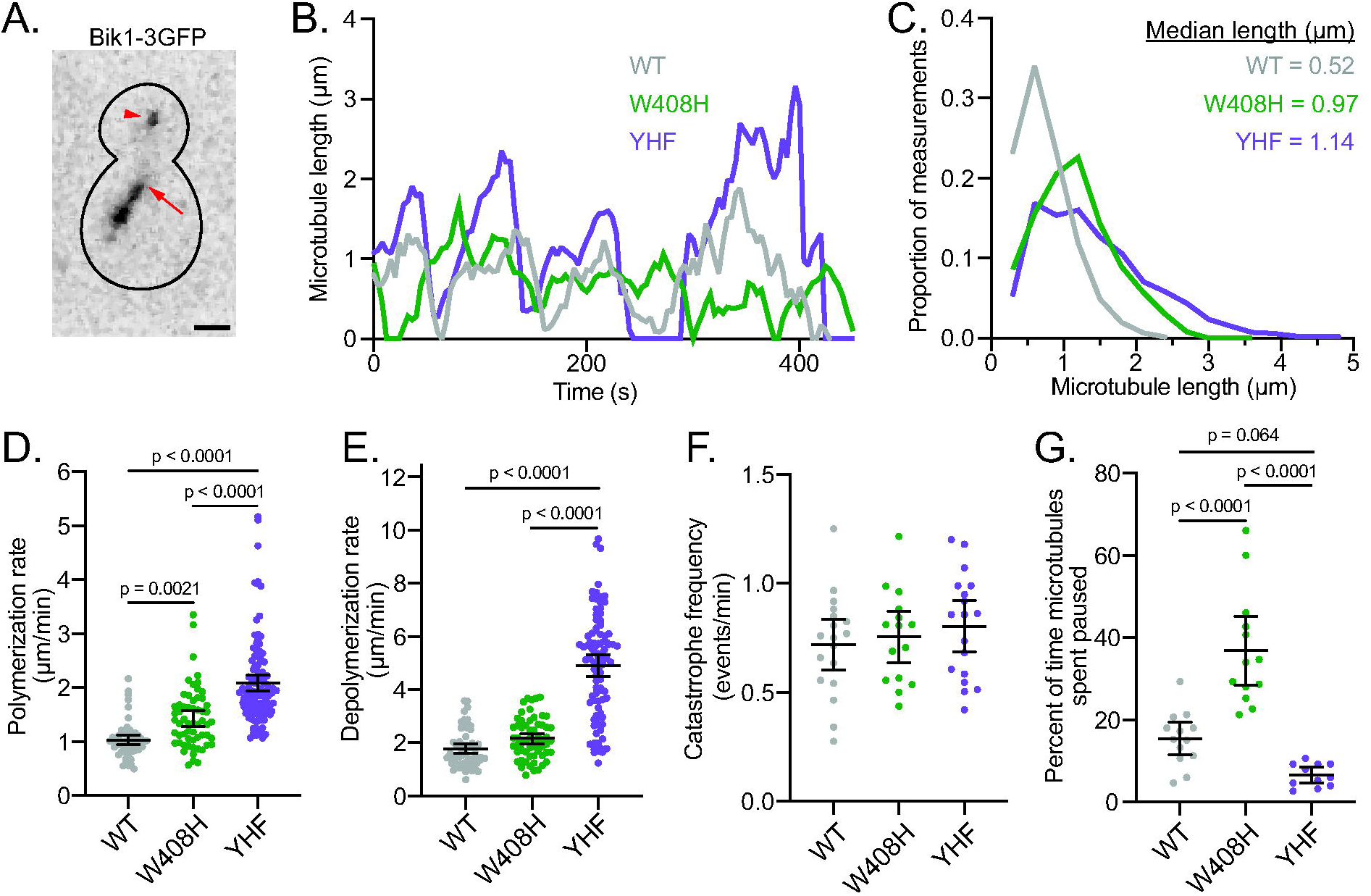
Naegleria α-tubulin helix 11’ variants alter microtubule dynamics. **(A)** Yeast cell expressing Bik1-3GFP. Arrowhead indicates plus end of astral microtubule while arrow indicates the proximal end of the mitotic spindle. Scalebar = 1 µm. **(B)** Lifeplots of individual astral microtubule length over time, with one representative microtubule each for wild type, *tub1*-W408H and *tub1*-YHF. **(C)** Distribution of all astral microtubule lengths measured. **(D)** Astral microtubule polymerization rates. Each dot represents a single polymerization event. **(E)** Astral microtubule depolymerization rate. Each dot represents an individual depolymerization event. **(F)** Castral microtubule catastrophe frequency. Each dot represents a catastrophe frequency from one cell imaged for 10 minutes. **(G)** Percent of time astral microtubules exhibit pause (see Materials and Methods). Each dot represents one cell imaged for 10 minutes. For panels D-G, bars and error bars represent represent mean value ± 95% confidence interval. Statistical analysis was done using an ordinary one-way ANOVA analyzed post hoc by a Tukey test.

Although both *tub1*-W408H and *tub1*-YHF mutants create long astral microtubules, they do so through distinct effects on microtubule dynamics. Cells expressing *tub1*-W408H exhibit significantly faster microtubule polymerization than wild-type cells (WT average: 1.0±0.09 µm/min, W408H average: 1.43±0.15; Figure 4D) but no significant difference in depolymerization (WT average: 1.77±0.18 µm/min, W408H average: 2.16±0.2; Figure 4E). In terms of transitions between polymerization and depolymerization, *tub1*-W408H cells do not exhibit a significant difference in catastrophe frequency (Figure 4F); however, we noticed that microtubules in these cells tend to maintain stable lengths for long periods of time. Maintenance of stable microtubule length over time has been described as the paused or attenuated state of microtubule dynamics^40,41,20^. To measure the pause state, we identified periods of time where microtubules neither assembled nor disassembled for at least 3 time points (8 seconds; see Materials and Methods). Whereas microtubules in wild-type control cells spend an average of 16% of their lifetimes in the pause state, microtubules in *tub1-*W408H cells spend 37% of their lifetime in pause (Figure 4G). We conclude that the W408H mutant increases microtubule polymerization and strongly promotes the pause state at the plus end.

In contrast, cells expressing *tub1*-YHF exhibit ∼2-fold faster polymerization (2.09±0.14 µm/min; Figure 4D) and depolymerization (4.89±0.41 µm/min; Figure 4E) than wild-type controls. Although *tub1*-YHF cells exhibit similar catastrophe frequency (Figure 4F), they spend less time in the pause state than *tub1-*W408H cells (Figure 4G). We conclude that the combination of F405Y, W408H, Y409F substitutions in the YHF mutant increases the exchange of heterodimers at the plus end, leading to faster polymerization, faster depolymerization and less time in pause. These results are consistent with our findings above that the F405Y, W408H, Y409F substitutions act synergistically to alter tubulin activity.

To reconcile the *tub1*-YHF triple mutant’s faster heterodimer exchange with the slower proliferation rate measured in Figure 3C, we examined how the mutant impacts mitotic spindle function. The primary role of microtubules in vegetatively growing budding yeast is to partition the duplicated genome, which requires the formation of a stable mitotic spindle with bi-oriented sister chromatids. Previous studies demonstrate that mutations in α– or β-tubulin that alter microtubule dynamics disrupt spindle formation, causing inefficient bi-orientation and prolonged mitosis via activation of the spindle assembly checkpoint (SAC;^42,43,44,45,46,47,48^. To determine whether the faster heterodimer exchange in *tub1*-YHF mutants disrupts spindle formation and leads to SAC arrest, we performed two experiments. First, we used live-cell imaging of mitotic cells to measure the separation distance between the Spindle Pole Bodies (SPBs; the budding yeast equivalent of centrosomes) over time. Budding yeast form a short spindle during S-phase and G2, that is maintained at a stable length until spindle elongation during anaphase (Figure 5A^49^). Consistent with this, we find that ‘pre-anaphase’ wild-type cells exhibit an average spindle length of 1.1 µm with a 23% coefficient of variation (Figure 5B-D). In contrast, *tub1*-YHF mutants exhibit shorter and less stable spindles, with an average length of 0.8 µm and 28% coefficient of variation (Figure 5B-D). This indicates that the F405Y, W408H, Y409F substitutions in the YHF mutant disrupts the formation of a stable spindle prior to anaphase. To test the prediction that *tub1-*YHF mutants require the SAC to prolong mitosis due to inefficient chromosome bi-orientation, we asked whether SAC genes are essential for the viability in *tub1-*YHF cells by using meiotic cross to create double mutants combining *tub1-*YHF with null mutations in the SAC genes *BUB1, BUB3* or *MAD2*. We were unable to recover viable double mutants combining *tub1-*YHF with *bub1*Δ or *bub3*Δ, indicating that these genes become essential in the presence of *tub1*-YHF. However, we did recover viable double mutants combining *tub1-*YHF and *mad2*Δ. In growth assays, these double mutants exhibit poor viability and greater sensitivity to benomyl than either *tub1-*YHF or *mad2*Δ single mutant alone (Figure 5E). This is consistent with a higher frequency of mitotic errors in *tub1-*YHF mutant cells that lack SAC activity. Together, these results indicate that the changes to heterodimer exchange at microtubule plus ends caused by the combination of F405Y, W408H, and Y409F substitutions in H11’ disrupts the formation of stable microtubule-kinetochore attachments.

**Figure 5.**
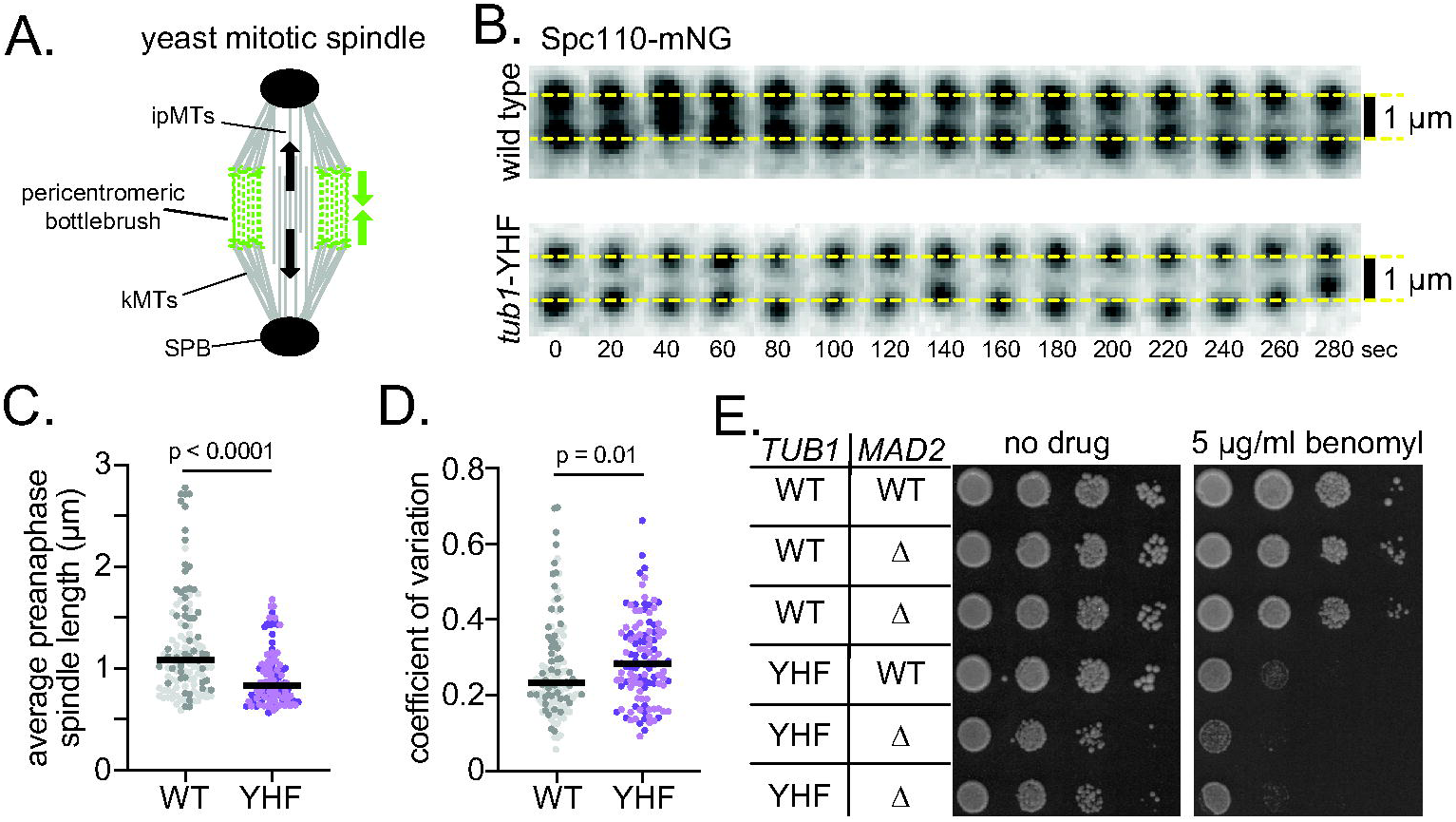
Naegleria α-tubulin helix 11’ variants destabilize mitotic spindles. **(A)** Schematic of the budding yeast mitotic spindle. ipMTs, interpolar microtubules; kMTs, kinetochore microtubules; SPB, Spindle Pole Body. The pericentromeric bottlebrush (green) describes the network of pericentromeric DNA loops extruded by cohesion and condensing that organize the budding yeast centromere and generate spring forces (green arrows) that counteract microtubule-based pulling forces (black arrows)^63^. **(B)** Sum intensity projections from time lapse Z-series images of Spc110-mNeonGreen labeled SPBs in a representative wild-type and *tub1-*YHF cell. Yellow dashed lines are separated by 1 µm and included for reference. **(C)** Average lengths of pre-anaphase spindle in wild-type and *tub1-*YHF cells. Each dot represents the average length from one cell imaged for 5 minutes. Colors indicate biological replicates. Bars represent median values of all per cell averages. Statistical analysis was done using the Mann-Whitney test. **(D)** Coefficient of variation for pre-anaphase spindle measurements. Each dot represents the average length from one cell imaged for 5 minutes. Colors indicate biological replicates. Bars represent median values of all per cell averages. Statistical analysis was done using the Mann-Whitney test. **(E)** Ten-fold dilution series of the indicated strains were spotted to rich medium (no drug) or rich medium supplemented with 5 µg/mL benomyl. Cells were grown at 30°C for two days.

### α-tubulin helix 11’ may support intradimer interactions with β-tubulin helix 8 in the curved heterodimer conformation

Finally, we sought to understand the role of H11’ in the structure and dynamics of the tubulin heterodimer. The transition of tubulin heterodimers from a curved to straight conformation is accompanied by reordering of interactions within the heterodimer^50^. We used computational modeling to determine how substitutions found in H11’ of *Naegleria* mitotic α-tubulins may impact important interactions in the curved or straight states of the heterodimer.

To test this, we first used the DynaMut2 server^51^ to visualize the interaction between α-tubulin H11’ residues and the surrounding regions in α– and β-tubulin in the curved heterodimer structure (PDB:6MZG;^28^), as well as a structure representing the straight state (PDB:3J7I;^52^). Both of these structures use tubulin heterodimers isolated from pig brain. DynaMut2 can currently only model single amino acid substitutions in a structure; therefore we chose to focus on Tryptophan 407 (W407, corresponding to W408 in yeast α-tubulin). When the heterodimer is in the curved conformation, we find that residue W407 is predicted to form hydrophobic, CH-π interactions with Alanine 254 (A254) in helix 8 of β-tubulin, (Figure 5A, first inset; hydrophobic bonds indicated by black dotted lines). When the heterodimer is in the straight conformation these bonds are absent, and W407 is instead predicted to form cation-π interactions with Arginine 251 (R251) and CH-π interactions with Valine 255 (V255) in β-tubulin helix 8 (Figure 5B, first inset). Using DynaMut2, we then changed α-tubulin W407 to Histidine in both structures, to mimic the variant residue in *Naegleria* mitotic α-tubulins. In the curved conformation, a histidine side chain at 407 is positioned further from β-tubulin helix 8 and is not predicted to interact with A254 in β-tubulin (Figure 5A, second inset). In the straight conformation, a histidine side chain at 407 maintains interactions with β-tubulin helix 8, but it is predicted to change the exact number and types of bonds. Specifically, Histidine 407 forms less extensive cation-π interactions with β-tubulin R251 and van der Waals with β-tubulin V255 (Figure 5B, second inset; van der Waals indicated by orange dotted line). These data suggest that the α-tubulin W407H substitution may destabilize the curved conformation while still stabilizing the straight conformation.

Our finding that W407 in α-tubulin H11’ may interact with β-tubulin helix 8 in the heterodimer, and that these interactions are altered for *Naegleria* α-tubulin, motivated us to ask whether β-tubulins in *Naegleria* exhibit variants in helix 8 that might accommodate this change. Similar to α-tubulin H11’, β-tubulin helix 8 is relatively well conserved across species (Figure 5C). By aligning β-tubulins from the same species described in Figure 2, we find that one of the two mitotic β-tubulins expressed in *Naegleria* exhibits a Serine substitution for Alanine at residue 254 (Figure 5C). This S254 residue was only observed in the *Naegleria* mitotic β-tubulin, and not the flagellate β-tubulins.

We next modeled the β-tubulin A254S variant using DynaMut2. In the curved heterodimer conformation, the A254S change in β-tubulin is predicted to form a hydrogen bond with α-tubulin W407 (Figure 5A, third inset). In the straight conformation, the A254S change in β-tubulin is not predicted to form any interactions with α-tubulin W407, and α-tubulin W407 is predicted to bind β-tubulin R251 and V255, similar to the wild-type structure (Figure 5B, first and third insets). Under current DynaMut2 parameters, we are unable to model α-tubulin W407H and β-tubulin A254S variants simultaneously in the same structure. Instead, we overlayed the α-W407H structure with the β-A254S structure to visualize how the two mutations may interact. Whereas the straight conformation of the merged α-W407H and β-A254S structure appears similar to the α-W407H model alone, the curved conformation shows the serine side chain of β-A254S positioned near the histidine side chain of α-W407H (Figure 5A-B, fourth insets). This close positioning could allow hydrogen bonding across the intradimer interface that is specific to the curved conformation. These results suggest that combining the β-tubulin helix 8 A254S substitution with the divergent α-tubulin H11’ may allow mitotic αβ-heterodimers to maintain intradimer interactions in both the curved and straight heterodimer states.

## Discussion

Our findings establish a role for α-tubulin H11’ in stabilizing the curved conformation of αβ-tubulin heterodimers and demonstrate the importance of this role for the exchange of heterodimers at microtubule plus ends. The strong conservation of H11’ across eukaryotes indicates that the conformational dynamics of the heterodimer is a core feature of tubulin evolution. Further, our results, together with our previous investigation of tubulinopathy missense variants at the V409 residue in H11’^23^, indicate that disrupting the conformational dynamics of the heterodimer may lead to developmental disease in humans. More broadly, our approach combining computational modeling of protein structure, comparative biology, and variant modeling in yeast provides a template for understanding the molecular impact of a growing number of missense variants in tubulinopathies.

Our findings for α-tubulin H11’ add to knowledge from previous studies of variants in α-and β-tubulin to highlight the importance of the conformational equilibrium of the heterodimer for microtubule dynamics. Biasing heterodimers toward a straight conformation is predicted to promote microtubule polymerization by restricting heterodimers in solution to a conformational state that is primed for interactions with neighboring tubulins in the microtubule lattice. Furthermore, straight heterodimers are predicted to inhibit catastrophe by preventing heterodimers in the lattice from returning to the curved conformation that weakens interactions with neighbors. Consistent with this, a tubulinopathy variant creating a D417H substitution in helix 12 of the β-tubulin isotype TUBB3 increases microtubule polymerization rate and decreases catastrophe frequency, and appears to bias the heterodimer toward a straight state based on weaker affinity for TOG domains and colchicine^21^. These results are similar to our previous findings for the V409A tubulinopathy variant in the α-tubulin isotype TUBA1A^23^. The *Naegleria* variants in α-tubulin H11’ in the current study also show increased polymerization, but do not appear to affect catastrophe (Figure 4). This suggests that the V409A and YHF variants have similar effects, but V409A may more strongly destabilize the curved conformation or strengthen the straight conformation. Conversely, biasing a heterodimer toward a curved conformation is predicted to slow microtubule polymerization and promote catastrophe by weakening interactions between heterodimers in the lattice. This is exemplified by the tubulinopathy variant R2W in the β-tubulin isotype TUBB4A, which decreases microtubule polymerization rate and increases catastrophes in vitro and in cells, and computational simulations predict that R2W shift heterodimers toward a curved conformation^22^. Together, these findings suggest that the conformational dynamics of heterodimers involve a coordination of interactions extending from the intradimer interface, to α-tubulin H11’ and β-tubulin helix 8, and extending to residues on β-tubulin helix 12. The identification of substitutions in these regions in tubulinopathy patients underscores the importance of conformational dynamics for tubulin function in vivo and suggests that examination of additional variants identified in tubulinopathy is likely to fill in the map of residues that contribute to these conformations. As the number of identified tubulinopathy variants continues to increase, open-access computational approaches such as those used here will be an important tool to prioritize variants for further study.

While H11’ is strongly conserved across eukaryotic α-tubulins, the conspicuous presence of variants in mitotic *Naegleria* α-tubulins suggests changes in evolutionary pressures on H11’ function. One possibility is that *Naegleria* mitosis may have evolved to favor heterodimers with limited conformational dynamics. To date, microtubule dynamics have not been measured in living *Naegleria*, however examination of mitotic spindles in fixed cells identified an unusual, barrel-shaped structure lacking focused spindle poles^35^. This raises the possibility of divergent spindle assembly mechanisms that could perhaps rely on straighter heterodimers that are more favorable for microtubule nucleation. A second possibility is that mitotic α-H11’ variants are accompanied by evolutionarily distinct MAPs with divergent mechanisms for regulating tubulin conformation and microtubule dynamics, as has been suggested by recent transcriptomic analysis of MAPs expressed in mitotic *Naegleria*^53^. A third possibility is that H11’ variants in *Naegleria* mitotic α-tubulins have co-evolved with changes in mitotic β-tubulins that together preserve the conformational dynamics of the heterodimer. Consistent with this, we find that a subset of mitotic β-tubulins in *Naegleria* exhibit a Serine at position 254 in helix 8, replacing the Alanine that is predicted to interact with αH11’ in the curved conformation (Figure 6). This suggests that the H11’ variants in mitotic α-tubulins either relieved selective pressure for A254 in β-tubulin, or vice versa, or that the substitution of Serine at position 254 creates a new function when combined with the H11’ variants. While we are unable to use DynaMut2 to assess the predicted changes in atomic interactions between α-tubulin W407H and β-tubulin A254S variants in combination, superimposing the predicted structures shows the longer Serine side chain at β-tubulin position 254 may compensate for the shorter Histidine side chain in α-tubulin position 407, and possibly form polar bonds when the heterodimer is in the curved conformation (Figure 6A). In contrast to human tubulinopathy, where de novo variants in α-tubulin H11’ create devastating disruption of microtubule function in neurons, the unique evolution of mitotic tubulins in *Naegleria* may have created an environment that is permissive or perhaps favorable to changes in H11’.

**Figure 6.**
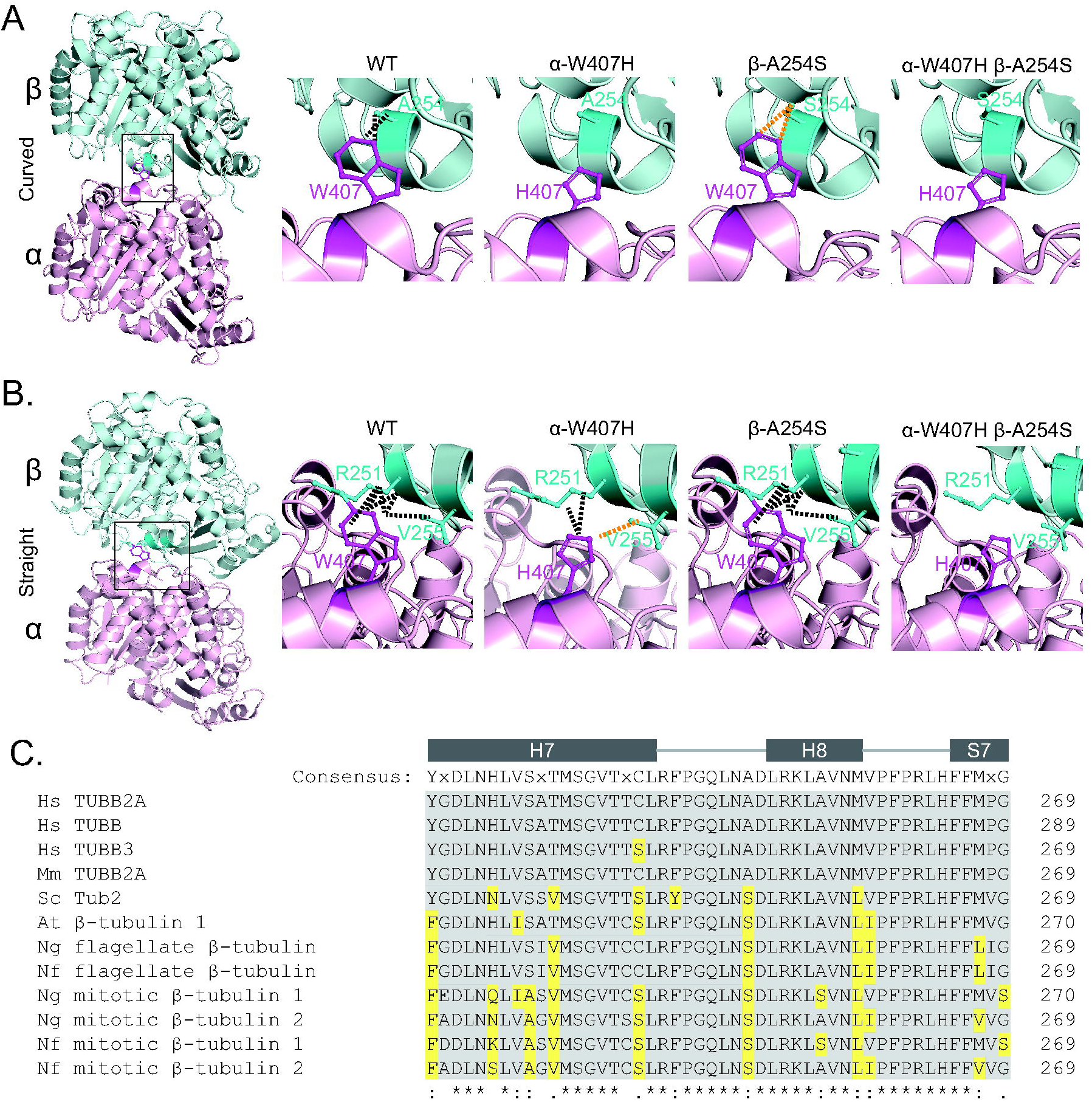
H11’ interacts with β-tubulin across the intradimer interface. **(A)**The DynaMut2 server was used to predict the binding interactions between α-tubulin residue 407 and residues in β-tubulin, using a model of the curved conformation of tubulin (PDB:6MZG^28^ Insets represent structures of either WT, *tub1*-W407H, *tub2*-A254S, or the overlay of the two mutants. Black and orange dotted lines represent predicted hydrophobic and polar interactions, respectively. **(B)** DynaMut2 predictions of binding interactions between α-tubulin residue 407 and residues in β-tubulin, using a model of the straight conformation of tubulin (PDB:3J7I)^52^. **(C)** Sequence alignment of β-tubulin helix 7, helix 8, and sheet 7 from a representative list of eukaryotes generated by Clustal Omega^55,56,57^. Consensus sequence generated from aligning 1,239 β-tubulins^34^. Grey and yellow backgrounds represent a conserved or divergent residue, respectively, when compared to the consensus sequence.

## Materials and Methods

### Protein conformation stability analysis using DUET

The web-based server DUET was used to estimate how mutations in α-tubulin impact the stability of the tubulin heterodimer structure based on predicted free energy changes^29^. The structure of *Sus scrofa* TUBA1A and TUBB α/β-tubulin heterodimer bound to the fission yeast XMAP215, Alp14 TOG1 and TOG2 domains (PDB:6MZG)^28^ was used to represent the curved heterodimer conformation. We instructed the DUET server to mutate the α-tubulin V409 residue to all other 19 amino acids and expanded this analysis on all residues in α-tubulin helix 11’. Plots in Figure 1B and 2D display the absolute value of the free energy changes of the imported protein structure that are predicted by DUET.

### Yeast strains and manipulations

All yeast experiments used *Saccharomyces cerevisiae* with the S288C genetic background. *TUB1*/α-tubulin mutants were generated by site-directed mutagenesis (Agilent Technologies; Sanata Clara, CA) of an integrating plasmid pJM469 containing a fragment of *TUB1* extending from bp 4 of the coding sequence to bp 487 3’ of the STOP codon. These plasmids were digested with XbaI and transformed into yeast, and integration at the native *TUB1* locus was confirmed by Sanger sequencing. Null mutants mad2Δ, bub1Δ, and bub3Δ were generated by PCR-mediated genome editing^54^. Plasmids, yeast strains, and oligos used in this study are listed in Tables S1, S2, and S3, respectively.

### Yeast growth assays

For doubling time assays in liquid media, yeast cells were grown to saturation before diluting 1:500 into fresh media, and 200µl of diluted cells were plated in each well of a 96-well plate. The OD_600_ values were measured every five minutes over the course of at least 19 hours on an Epoch 2 Microplate Spectrophotometer (BioTek #EPOCH2NS; Winooski, VT) that was incubated at 30°C with continuous orbital shaking. The OD_600_ values were then fitted to a logistic curve using Graphpad Prism, and doubling time was calculated by dividing the natural log of 2 by the rate constant (k). Statistical analyses were conducted on GraphPad Prism using an ordinary one-way ANOVA and corrected for multiple comparisons post hoc by a Tukey test. For proliferation assays in the presence of nocodazole, the DMSO control and 1.25 µM nocodazole data for each genotype were compared using the Mann-Whitney test in GraphPad Prism.

For proliferation assays on solid media, measuring sensitivity to benomyl and low temperature, yeast cells were grown to saturation in rich media and a 10-fold dilution series was created in a 96-well plate. Suspensions of diluted cells were spotted to either rich media plates or rich media plates containing 5, 10, or 15µg/ml benomyl. Plates were grown at the time and temperature indicated in the figure legends.

### Sequence alignments

1,347 α-tubulin sequences from 438 species^34^ were aligned using the Multiple Sequence Alignment from Clustal Omega^55,56,57^. The sequence alignment was then downloaded, saved as a plain text file, and viewed with SnapGene software (www.snapgene.com), which also presented the consensus sequence calculated from this multiple sequence alignment.

### Conservation analysis using ConSurf

The α-tubulin multiple sequence alignment described above was used to determine the conservation of each amino acid in the consensus sequence using the ConSurf server^36,37^. The conservation score was mapped onto the imported structure of *Sus scrofa* TUBA1A and TUBB (PDB:6MZG)^28^ using the ConSurf protein structure feature^58,59^. The mean and standard deviations of the percentages of α-tubulins that maintain the same amino acid as found in the consensus sequence for each TUBA1A secondary structure, as seen in Table 1, were also calculated from ConSurf analysis.

### Imaging GFP-Tub1 in budding yeast

Cells expressing either WT *TUB1*, *tub1*-F405Y, *tub1*-W408H, *tub1*-Y409F, or *tub1*-F405Y-W408H-Y409F at the endogenous locus were transformed with plasmids with GFP-Tub1, GFP-*tub1*-F405Y, GFP-*tub1*-W408H, GFP-*tub1*-Y409F, or GFP-*tub1*-F405Y-W408H-Y409F, respectively. Mutant plasmids were generated by site-directed mutagenesis of the GFP-Tub1 integration plasmid (pSK1050)^60^. Transformed cells were grown overnight at 30°C in selection media, then diluted the following morning into fresh imaging media and incubated while shaking until they were in log-phase. Cells were imaged on a Nikon Ti-E spinning disk confocal (CSU10; Yokogawa) equipped with a 1.45 NA 100x CFI Plan Apo oil objective, a piezo electric stage (Physik Instrumente, Auburn, MA), a 488 nm laser, and an EMCCD camera (iXon Ultra 897; Andor Technology, Belfast, UK). Images were collected using the NIS Elements software (Nikon). Z-stack steps of 0.4µm were taken over a range of 7.2µm at 30°C. At least 30 cells were analyzed per genotype. Representative images shown are sum projections of these Z-stacks. Scale bars represent 1µm.

### Yeast microtubule dynamics

Cells expressing Bik1-3GFP with either WT *TUB1*, *tub1*-F405Y, *tub1*-W408H, *tub1*-Y409F, or *tub1*-F405Y-W408H-Y409F were grown to log-phase at 30°C in non-fluorescent media. Imaging slide chambers were prepared with concanavalin A to adhere cells to the slide. Chambers were sealed with VALAP (Vaseline, lanolin, and paraffin at a 1:1:1 ratio). Cells were imaged on the same Nikon Ti-E spinning disk confocal described above. Z-series steps of 0.4µm over a 7.2µm range were imaged every four seconds for 10 minutes at 30°C. To measure microtubule dynamics, astral microtubule lengths in pre-anaphase cells were measured at each time point and lengths were defined as the distance from the spindle pole body to the microtubule plus end. Microtubule dynamics were measured using a previously described MATLAB code^61^. Paused microtubules were defined as periods where microtubules exhibit less than 500nm of length change across a time period of at least 3 consecutive time points^20^. All cell genotypes were blinded for microtubule measurements and analysis. At least 24 cells were analyzed for all genotypes. Statistical analyses were performed for each test using an ordinary one-way ANOVA and corrected for multiple comparisons by using a Tukey test.

### Pre-anaphase spindle stability

Cells expressing Spc110-mNeonGreen were grown in rich liquid media at 30°C, then diluted and grown to log phase in fresh, synthetic media. Cells were imaged at 30°C on spinning disk confocal. Z-series consisting of a 6 µm range at 0.4 µm steps were acquired every 20 seconds for 5 minutes. Pre-anaphase cells were identified in image series of asynchronous cultures based on bud size and spindle length. Spindle length was defined as the linear distance in 3 dimensions between the centroid of the Spc110-mNeonGreen signal, and measured for every Z-series at each point in the time course. Average spindle lengths and coefficient of variation calculated for each cell analyzed were compared using the Mann-Whitney test in GraphPad Prism.

### Modeling predicted atomic interactions using DynaMut2

Predictive atomic interactions between residues of interest were done using the online server DynaMut2^51^. Two structures were used in this analysis: PDB:6MZG^28^ to represent the curved state of tubulin and PDB:3J7I^52^ to represent the straight state of tubulin. Predicted structures and interactions for WT and mutant states were downloaded as PDB files and visualized using PyMOL (The PyMOL Molecular Graphics System, Version 2.0 Schrödinger, LLC).

## Acknowledgements

We are grateful to members of the Moore lab for helpful discussions. This work was supported by Bolie Scholar Award (K.J.H) and NIH R35GM136253 (J.K.M.).

