## Supplemental Table 1 for "Role of α-tubulin helix 11’ in heterodimer conformation and microtubule dynamics"

**Table S1. Yeast strains used in this study**

| **Yeast Strain** | **Genotype** | **Source** |
| --- | --- | --- |
| 1840 | *MATα ura3-52 lys2-801 leu2-∆1 his3-∆200 trp1-∆63* | This study |
| 3711 | *MATα tub1::pJM507(TUB1-V410I +487::URA) ura3-52 lys2-801 leu2-∆1 his3-∆200 trp1-∆63* | This study |
| 3768 | *MATA tub1::pJM560(TUB1-V410A +487::URA) ura3-52 lys2-801 leu2-∆1 his3-∆200 trp1-∆63* | This study |
| 3769 | *MATα tub1::pJM560(TUB1-V410A +487::URA) ura3-52 lys2-801 leu2-∆1 his3-∆200 trp1-∆63* | This study |
| 3957 | *MATα tub1::pJM560(TUB1-V410I +487::URA) ura3-52 lys2-801 leu2-∆1 his3-∆200 trp1-∆63* | This study |
| 4918 | *MATA tub1::pJM769(TUB1-V410D +487::URA) ura3-52 lys2-801 leu2-∆1 his3-∆200 trp1-∆63* | This study |
| 4919 | *MATα tub1::pJM769(TUB1-V410D +487::URA) ura3-52 lys2-801 leu2-∆1 his3-∆200 trp1-∆63* | This study |
| 5193 | *MATA tub1::pJM853(TUB1-F405Y +487::URA) ura3-52 lys2-801 leu2-∆1 his3-∆200 trp1-∆63* | This study |
| 5194 | *MATα tub1::pJM853(TUB1-F405Y +487::URA) ura3-52 lys2-801 leu2-∆1 his3-∆200 trp1-∆63* | This study |
| 5198 | *MATα tub1::pJM854(TUB1-F405Y-W408H-Y409F +487::URA) ura3-52 lys2-801 leu2-∆1 his3-∆200 trp1-∆63* | This study |
| 5199 | *MATα tub1::pJM854(TUB1-F405Y-W408H-Y409F +487::URA) ura3-52 lys2-801 leu2-∆1 his3-∆200 trp1-∆63* | This study |
| 5201 | *MATα tub1::pJM855(TUB1-W408H +487::URA) ura3-52 lys2-801 leu2-∆1 his3-∆200 trp1-∆63* | This study |
| 5202 | *MATA tub1::pJM855(TUB1-W408H +487::URA) ura3-52 lys2-801 leu2-∆1 his3-∆200 trp1-∆63* | This study |
| 5205 | *MATα tub1::pJM856(TUB1-Y409F +487::URA) ura3-52 lys2-801 leu2-∆1 his3-∆200 trp1-∆63* | This study |
| 5206 | *MATA tub1::pJM856(TUB1-Y409F +487::URA) ura3-52 lys2-801 leu2-∆1 his3-∆200 trp1-∆63* | This study |
| 5222 | *MATα tub1::pJM469(TUB1 +487::URA) ura3-52 lys2-801 leu2-∆1 his3-∆200 trp1-∆63* | This study |
| 5223 | *MATA tub1::pJM469(TUB1 +487::URA) ura3-52 lys2-801 leu2-∆1 his3-∆200 trp1-∆63* | This study |
| 5290 | *MATα tub1::pJM469(TUB1 +487::URA) BIK1-3GFP::TRP1 ura3-52 lys2-801 leu2-∆1 his3-∆200 trp1-∆63* | This study |
| 5291 | *MATA tub1::pJM469(TUB1 +487::URA) BIK1-3GFP::TRP1 ura3-52 lys2-801 leu2-∆1 his3-∆200 trp1-∆63* | This study |
| 5294 | *MATA tub1::pJM855(TUB1-W408H +487::URA) BIK1-3GFP::TRP1 ura3-52 lys2-801 leu2-∆1 his3-∆200 trp1-∆63* | This study |
| 5295 | *MATα tub1::pJM855(TUB1-W408H +487::URA) BIK1-3GFP::TRP1 ura3-52 lys2-801 leu2-∆1 his3-∆200 trp1-∆63* | This study |
| 5298 | *MATα tub1::pJM854(TUB1-F405Y-W408H-Y409F +487::URA) BIK1-3GFP::TRP1 ura3-52 lys2-801 leu2-∆1 his3-∆200 trp1-∆63* | This study |
| 5299 | *MATΑ tub1::pJM854(TUB1-F405Y-W408H-Y409F +487::URA) BIK1-3GFP::TRP1 ura3-52 lys2-801 leu2-∆1 his3-∆200 trp1-∆63* | This study |
| 5570 | *MATα tub1::pJM469(TUB1 +487::URA) Spc110-mNeonGreen::HIS3 ura3-52 lys2-801 leu2-∆1 his3-∆200 trp1-∆63* | This study |
| 5571 | *MATα tub1::pJM469(TUB1 +487::URA) Spc110-mNeonGreen::HIS3 ura3-52 lys2-801 leu2-∆1 his3-∆200 trp1-∆63* | This study |
| 5572 | *MATΑ tub1::pJM854(TUB1-F405Y-W408H-Y409F +487::URA) Spc110-mNeonGreen::HIS3 ura3-52 lys2-801 leu2-∆1 his3-∆200 trp1-∆63* | This study |
| 5573 | *MATα tub1::pJM854(TUB1-F405Y-W408H-Y409F +487::URA) Spc110-mNeonGreen::HIS3 ura3-52 lys2-801 leu2-∆1 his3-∆200 trp1-∆63* | This study |
| 5602 | *MATA tub1::pJM854(TUB1-F405Y-W408H-Y409F +487::URA) mad2∆::HIS3 ura3-52 lys2-801 leu2-∆1 his3-∆200 trp1-∆63* | This study |
| 5603 | *MATA tub1::pJM854(TUB1-F405Y-W408H-Y409F +487::URA) mad2∆::HIS3 ura3-52 lys2-801 leu2-∆1 his3-∆200 trp1-∆63* | This study |
| 5604 | *MATA tub1::pJM854(TUB1 +487::URA) mad2∆::HIS3 ura3-52 lys2-801 leu2-∆1 his3-∆200 trp1-∆63* | This study |
| 5605 | *MATA tub1::pJM854(TUB1 +487::URA) mad2∆::HIS3 ura3-52 lys2-801 leu2-∆1 his3-∆200 trp1-∆63* | This study |
| 5903 | *MATA tub1::pJM853(TUB1-F405Y +487::URA) PDR1::pdr1-DBD-CYC8::LEU2 ura3-52 lys2-801 leu2-∆1 his3-∆200 trp1-∆63* | This study |
| 5904 | *MATA tub1::pJM853(TUB1-F405Y +487::URA) PDR1::pdr1-DBD-CYC8::LEU2 ura3-52 lys2-801 leu2-∆1 his3-∆200 trp1-∆63* | This study |
| 5905 | *MATA tub1::pJM854(TUB1-F405Y-W408H-Y409F +487::URA) PDR1::pdr1-DBD-CYC8::LEU2 ura3-52 lys2-801 leu2-∆1 his3-∆200 trp1-∆63* | This study |
| 5906 | *MATα tub1::pJM854(TUB1-F405Y-W408H-Y409F +487::URA) PDR1::pdr1-DBD-CYC8::LEU2 ura3-52 lys2-801 leu2-∆1 his3-∆200 trp1-∆63* | This study |
| 5907 | *MATα tub1::pJM854(TUB1-W408H +487::URA) PDR1::pdr1-DBD-CYC8::LEU2 ura3-52 lys2-801 leu2-∆1 his3-∆200 trp1-∆63* | This study |
| 5908 | *MATα tub1::pJM854(TUB1-W408H +487::URA) PDR1::pdr1-DBD-CYC8::LEU2 ura3-52 lys2-801 leu2-∆1 his3-∆200 trp1-∆63* | This study |
| 5909 | *MATα tub1::pJM856(TUB1-Y409F +487::URA) PDR1::pdr1-DBD-CYC8::LEU2 ura3-52 lys2-801 leu2-∆1 his3-∆200 trp1-∆63* | This study |
| 5910 | *MATα tub1::pJM856(TUB1-Y409F +487::URA) PDR1::pdr1-DBD-CYC8::LEU2 ura3-52 lys2-801 leu2-∆1 his3-∆200 trp1-∆63* | This study |
| 5911 | *MATα tub1::pJM469(TUB1 +487::URA) PDR1::pdr1-DBD-CYC8::LEU2 ura3-52 lys2-801 leu2-∆1 his3-∆200 trp1-∆63* | This study |
| 5912 | *MATα tub1::pJM469(TUB1 +487::URA) PDR1::pdr1-DBD-CYC8::LEU2 ura3-52 lys2-801 leu2-∆1 his3-∆200 trp1-∆63* | This study |
| 5913 | *MATA tub1::pJM469(TUB1 +487::URA) PDR1::pdr1-DBD-CYC8::LEU2 ura3-52 lys2-801 leu2-∆1 his3-∆200 trp1-∆63* | This study |
