## Supplemental Table 2 for "Role of α-tubulin helix 11’ in heterodimer conformation and microtubule dynamics"

**Table S2. Plasmids used in this study**

| **Plasmid ID** | **Plasmid name** | **Source** |
| --- | --- | --- |
| pJM469 | TUB1-pRS306 integrating plasmid | Aiken et al. 2020 |
| pJM507 | tub1-V410I-pRS306 integrating plasmid | Hoff et al. 2022 |
| pJM560 | tub1-V410A-pRS306 integrating plasmid | Hoff et al. 2022 |
| pJM769 | tub1-V410D-pRS306 integrating plasmid | This study |
| pJM853 | tub1-F405Y-pRS306 integrating plasmid | This study |
| pJM854 | tub1-F405Y-W408H-Y409F-pRS306 integrating plasmid | This study |
| pJM855 | tub1-W408H-pRS306 integrating plasmid | This study |
| pJM856 | tub1-Y409F-pRS306 integrating plasmid | This study |
