## Supplemental Table 3 for "Role of α-tubulin helix 11’ in heterodimer conformation and microtubule dynamics"

**Table S3. Oligonucleotides used in this study**

| **Oligo ID** | **Oligo name** | **Oligo sequence** | **Source** |
| --- | --- | --- | --- |
| 545 | Tub1-V410A F | CGTCCACTGGTATGcCGGTGAAGGTATG | Hoff et al. 2022 |
| 546 | Tub1-V410A R | CATACCTTCACCGgCATACCAGTGGACG | Hoff et al. 2022 |
| 1018 | Tub1-V410I F | CGTGCTTTCGTCCACTGGTATaTCGGTGAAGGT | Hoff et al. 2022 |
| 1019 | Tub1-V410I R | ACCTTCACCGAtATACCAGTGGACGAAAGCACG | Hoff et al. 2022 |
| 1613 | Tub1-V410D F | TTTCGTCCACTGGTATGaCGGTGAAGGTATGGAAG | This study |
| 1614 | Tub1-V410D R | CTTCCATACCTTCACCGtCATACCAGTGGACGAAA | This study |
| 1861 | Tub1-F405Y F | AATGTATGCCAAACGTGCTTatGTCCACTGGTATGTCGGT | This study |
| 1862 | Tub1-F405Y R | ACCGACATACCAGTGGACatAAGCACGTTTGGCATACATT | This study |
| 1863 | Tub1-F405Y_W408H_Y409F F | CGATAGAAAATTCGATTTAATGTATGCCAAACGTGCTTatGTCCACcatTtTGTCGGTGAAGGTATGGAAG | This study |
| 1864 | Tub1-F405Y_W408H_Y409F R | CTTCCATACCTTCACCGACAaAatgGTGGACatAAGCACGTTTGGCATACATTAAATCGAATTTTCTATCG | This study |
| 1727 | Tub1-W408H F | GCCAAACGTGCTTTCGTCCACcatTATGTCGGTGAAGGTATGGAA | This study |
| 1728 | Tub1-W408H R | TTCCATACCTTCACCGACATAatgGTGGACGAAAGCACGTTTGGC | This study |
| 1731 | Tub1-Y409F F | GTGCTTTCGTCCACTGGTtTGTCGGTGAAGGTATG | This study |
| 1732 | Tub1-Y409F R | CATACCTTCACCGACAaACCAGTGGACGAAAGCAC | This study |
